# Kinase activity of histone chaperone APLF maintains steady state of centrosomes in mouse embryonic stem cells

**DOI:** 10.1101/2024.02.06.579158

**Authors:** Sruthy Manuraj Rajam, Pallavi Chinnu Varghese, Mayur Balkrishna Shirude, Khaja Mohieddin Syed, Anjali Devarajan, Kathiresan Natarajan, Debasree Dutta

**Affiliations:** Rajiv Gandhi Centre for Biotechnology (RGCB), Regenerative Biology Program, Thycaud PO, Poojappura, Thiruvananthapuram 695014, Kerala, India; Manipal Academy of Higher Education, Manipal, Karnataka State, 576104, India; Department of Molecular and Cell Biology, University of California, Berkeley, CA 94720-3370, USA; Rajiv Gandhi Centre for Biotechnology (RGCB), Transdisciplinary Biology Program, Thycaud PO, Poojappura, Thiruvananthapuram 695014, Kerala, India

**Keywords:** APLF, Histone chaperone, DNA repair, centrosome, PLK4, kinase

## Abstract

Our recent studies revealed the role of mouse Aprataxin PNK-like Factor (APLF) in development. Nevertheless, the comprehensive characterization of mouse APLF remains entirely unexplored. Based on domain deletion studies, here we report that mouse APLF’s Acidic Domain and Fork Head Associated (FHA) domain can chaperone histones and repair DNA like the respective human orthologs. Immunofluorescence studies in mouse embryonic stem cells showed APLF co-localized with γ-tubulin within and around the centrosomes and govern the number and integrity of centrosomes via PLK4 phosphorylation. Enzymatic analysis established mouse APLF as a kinase. Docking studies identified three putative ATP binding sites within the FHA domain. Site-directed mutagenesis showed that R37 residue is indispensable for the kinase activity of APLF thereby regulating the centrosome number. These findings might assist us comprehend APLF in different pathological and developmental conditions and reveal non-canonical kinase activity of proteins harbouring FHA domains that might impact multiple cellular processes.

## Introduction

Histone chaperones are histone escort proteins that regulate chromatin accessibility for DNA-templated processes like DNA repair, gene regulation, and replication. Aprataxin Polynucleotide Kinase-like Factor (APLF) is DNA damage dependent histone chaperone responsible for the recruitment and eviction of H2A-H2B/H3-H4 from the nucleosome (Mehrotra et al., 2011; Corbeski et al., 34 2018; Corbeski et al., 2022). Human APLF is a multidomain protein that encompasses five functional domains. The N-terminus Forkhead Association Domain (FHA) along with Ku binding motif (KBM) and tandem PAR-Binding Zinc Finger Domain (PBZ) are responsible for the Non-Homologous End Joining (NHEJ) repair whereas the C-terminus Acidic Domain (AD) harbors the histone chaperone activity (Mehrotra et al., 2011; Corbeski et al., 2018; Corbeski et al., 2022). APLF^AD^ contains the Histone binding Domain (HBD) (Corbeski et al., 2018). APLF, with its FHA domain and KBM, serves as a scaffold for the NHEJ complex to assemble and facilitate the DNA end processing at the damage site (Macrae et al.,2008; Shirodkar et al., 2013). Human APLF has been extensively studied in the progression of diseases including breast cancer, bladder cancer, Glioblastoma multiforme (Majumder et al., 2018; Dong W et al.,2020; Ritcher et al., 2019). Our earlier study showed enhanced expression of APLF promotes metastasis-associated Epithelial-Mesenchymal transition (EMT) in breast cancer by deregulating the recruitment of macroH2A.1 to the EMT-specific gene promoters (Majumder et al., 2018). Several studies in human APLF explain its structure-function relation. However, the studies in its mouse ortholog are very limited. Nonetheless, we recently reported the function of murine APLF in the generation of induced pluripotency in murine fibroblast (Syed et al., 2016) and implantation of the mouse embryo (Varghese et al., 2021). These findings collectively describe APLF as a factor that has profound implications in cells with high division potential. Hence, it is of great importance to characterize murine APLF functionally.

In this study, we employed mouse ESCs as the experimental model to examine the functional properties of mouse APLF in facilitating cellular mechanisms. The investigation on domain-specific and localization-specific functioning has converged on the identification of a previously unknown characteristic of murine APLF or in that case, for any other histone chaperone. This mechanistic insight might improve our understanding of cellular processes mediated by APLF or proteins possessing a FHA domain.

## Results

### Murine APLF domains share similar function to their human orthologs

In 2016, for the first time we reported on the role of murine APLF in induction of pluripotency (Syed et al., 2016). Interestingly, human and mouse APLF proteins display only ∼59% similarity in the amino acid sequences (Fig. 1A). However, pairwise alignment of amino acids sequence showed the presence of similar functional domains in mouse compared to the human APLF protein (Fig. S1). Human APLF has two PAR-binding domains (PBZ), a ku binding domain, a Fork Head Association (FHA) domain and a C-terminal Acidic domain (AD) (Fig. S1). Pairwise sequence alignment indicated the presence of all these domains in mouse APLF (Fig. S1). Although we have reported on the various roles of mouse APLF in pluripotency or pre- implantation development of embryo, experimental evidences on the function of the domains has not yet been elucidated till date. NHEJ mediated DNA repair and histone chaperone activity are the two major functions of APLF. Based on pairwise alignment of human and mouse APLF protein sequences, we generated GFP tagged deletion constructs for N terminus FHA domain (ΔFHA) and C terminus Acidic domain (ΔAD), which might abrogate the DNA repair and histone chaperone activity respectively (Fig. 1B). Full length *Aplf* cDNA was PCR amplified from mouse embryonic fibroblasts (MEFs). The reconstitution of the nucleosome is the fundamental function of a histone chaperone. Hence, to study its function *in vivo*, the ability of histone chaperones to recruit and replenish the depleted histones in the nucleus from its pool, can be assessed by the recovery of fluorescence in the bleached nucleus or by Fluorescence recovery after photobleaching (FRAP) assay. HEK293T cells were co-transfected with GFP-tagged *Aplf* domain clones (Fig. 1C) with H2B-mCherry (Addgene, #20972) (Nam and Benezra, 2009). Representative confocal images demonstrate pre-bleaching, bleaching and post-bleaching recovery of histone H2B in nucleus of cells transfected with APLF domain-deletion constructs. (Fig. 1D). Significant increase in the half time for recovery after photobleaching was observed in cells transfected with ΔAD compared to other constructs (Fig. 1E) whereas a significant decrease in mobile fraction was observed in the absence of AD (Fig. 1F). This substantiated the fact that the AD of APLF could replenish the depleted H2B to the bleached area in the nucleus, thereby confirming its histone chaperone activity. To assess the DNA repair capability of different domains of APLF, HEK293T cells upon transfection with the same constructs, were treated with 10μM etoposide for 4hours to induce DNA double strand (DSB) breaks or DNA damage followed by their recovery for 24hours (Syed et al., 2016). The rapid and ubiquitous phosphorylation of H2AX residue Ser139 (γH2AX) after DNA DSB formation has been extensively explored in APLF and NHEJ contexts (Fenton et al., 2013; Tong et al., 2016). Interestingly loss in the FHA domain resulted in significant retention of the γH2A.X foci indicating compromise of the DNA repair activity in absence of the domain (Fig. 1G, S2). Collectively, these findings validate the domain specific functional conservation of murine APLF with human APLF as DNA repair factor and histone chaperone with its FHA and Acidic domains respectively.

**Figure 1.**
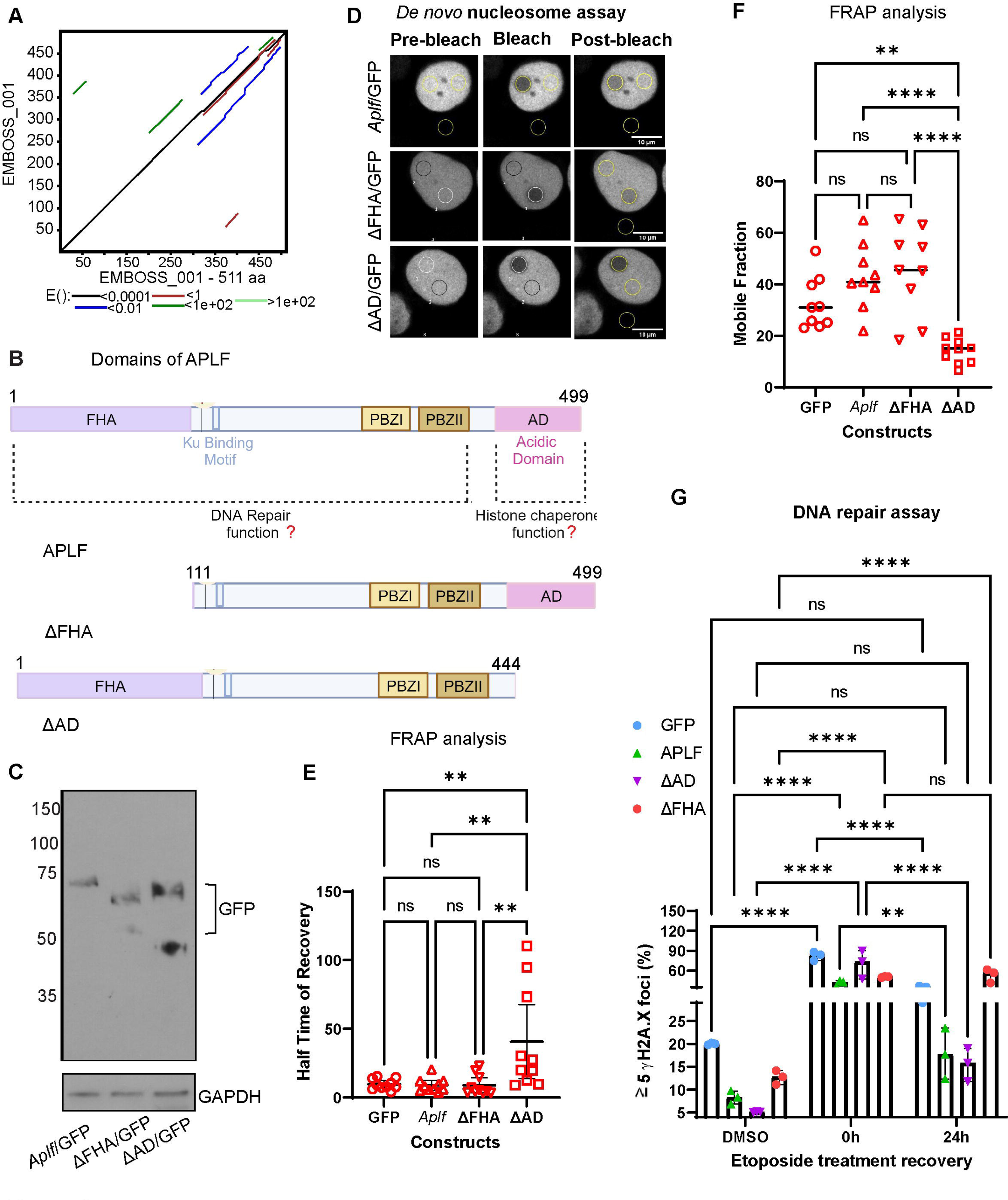
Mouse APLF domains exhibit histone chaperoning and DNA repair activity. A. Plot representing the pairwise sequence alignment of amino acids in mouse vs. human APLF proteins using LALIGN bioinformatics tool (https://www.ebi.ac.uk/Tools/services/web/toolresult.ebi?jobId=lalign-I20231015-184430-0525-54532945-p1m&context=protein&analysis=visual) (Madeira et al., 2022). Pairwise sequence alignment was generated by EMBOSS tool, wherein red represent small and hydrophobic, including aromatic amino acids; blue represent acidic, magenta represent basic and green represent hydroxyl/sulfhydryl/amine/Glycine residues. B. Domain specific constructs of mouse APLF, based on alignment with human APLF (refer to Fig. S1). C. Western blot analysis for the expression of GFP-tagged APLF constructs. Individual APLF domains were cloned in pEGFPN1 vector and transfected in HEK293T cells. Protein was isolated in RIPA buffer from HEK293T cells and run on SDS-PAGE and probed with GFP antibody. D-F. FRAP analysis for the detection of *in vivo* nucleosome assay mediated by APLF. GFP-tagged APLF constructs expressing HEK 293T cells were co-transfected with H2B-mCherry plasmid and subjected to FRAP analysis. Photobleaching was done at ROI in H2B:mCherry expressing cells and were monitored for the fluorescence recovery in the bleached region (D). Graphical representation for halftime of recovery, E, and mobile fraction present, F, in cells harboring different constructs of APLF. Ordinary one-way ANOVA statistical analysis was done with Tukey’s multiple comparisons tests, ****p<0.0001. G. HEK293T cells transfected with GFP tagged APLF domain constructs were subjected to 10mM etoposide or DMSO for 4 hours to induce double strand DNA damage followed by its release and observation for the formation of gH2A.X foci by immunofluorescence analysis at 0 and 24 hours of its release to monitor the DNA repair activity. Graphical representation of the DNA repair assay mediated in presence of APLF full length, ΔFHA, ΔAD clones and control cells. A two-way ANOVA statistical analysis was done followed by Tukey’s multiple comparisons tests, ****p<0.0001.

But, which interacting partners associate with mouse APLF in mediating these functions?

### APLF interactome reveals novel interaction partners

To further characterize the protein APLF, we generated FLAG-tagged APLF clones (full length, ΔAD and ΔFHA domains) and transfected into HEK293T cells (Fig. S3A). Subsequently proteins isolated were subjected to LC/MS-MS analysis. The examination of the STRING database (Szklarczyk et al., 2023) revealed the presence of seven distinct clusters of interaction partners associated with APLF (Fig. 2A). Mouse APLF interacted with components of the nucleosome assembly, double strand DNA repair, RNA metabolism, metabolic processes, protein folding, cytoplasmic translation and cytoskeletal organization (Fig. 2A). Gene ontology (GO) analysis for top ten biological processes indicated APLF associated with regulation of centromere assembly complex, mRNA splicing/spliceosome along with obvious ones of nucleosome assembly and DNA repair (Fig. 2B). GO analysis for the cellular component (Fig. 2C) showed APLF being present within the cytoplasm, CENPA-containing nucleosome and cytoskeleton along with earlier known cellular compartments including nucleus, nucleosome among others. Next, we determined the interacting partners of APLF-ΔAD and APLF-ΔFHA domain constructs. STRING database analysis indicated the clustering of ΔFHA domain resulted in nucleosome organization, cytoplasmic proteins and cytoskeleton organization but not the microtubule organization (Fig. 2D), unlike its presence in ΔAD associated clusters. STRING database analysis indicated the clustering of ΔAD domain with all those common to full length APLF except for the nucleosome assembly cluster (Fig. 2E). Thus, the study on the APLF interactome not only confirmed the role of mouse APLF in DNA repair and histone chaperone activity, but also suggested a potential involvement of the FHA domain of APLF in the regulation of cytoskeletal events such as microtubule organization or centrosome organization, which indicate a novel function of APLF.

**Figure 2.**
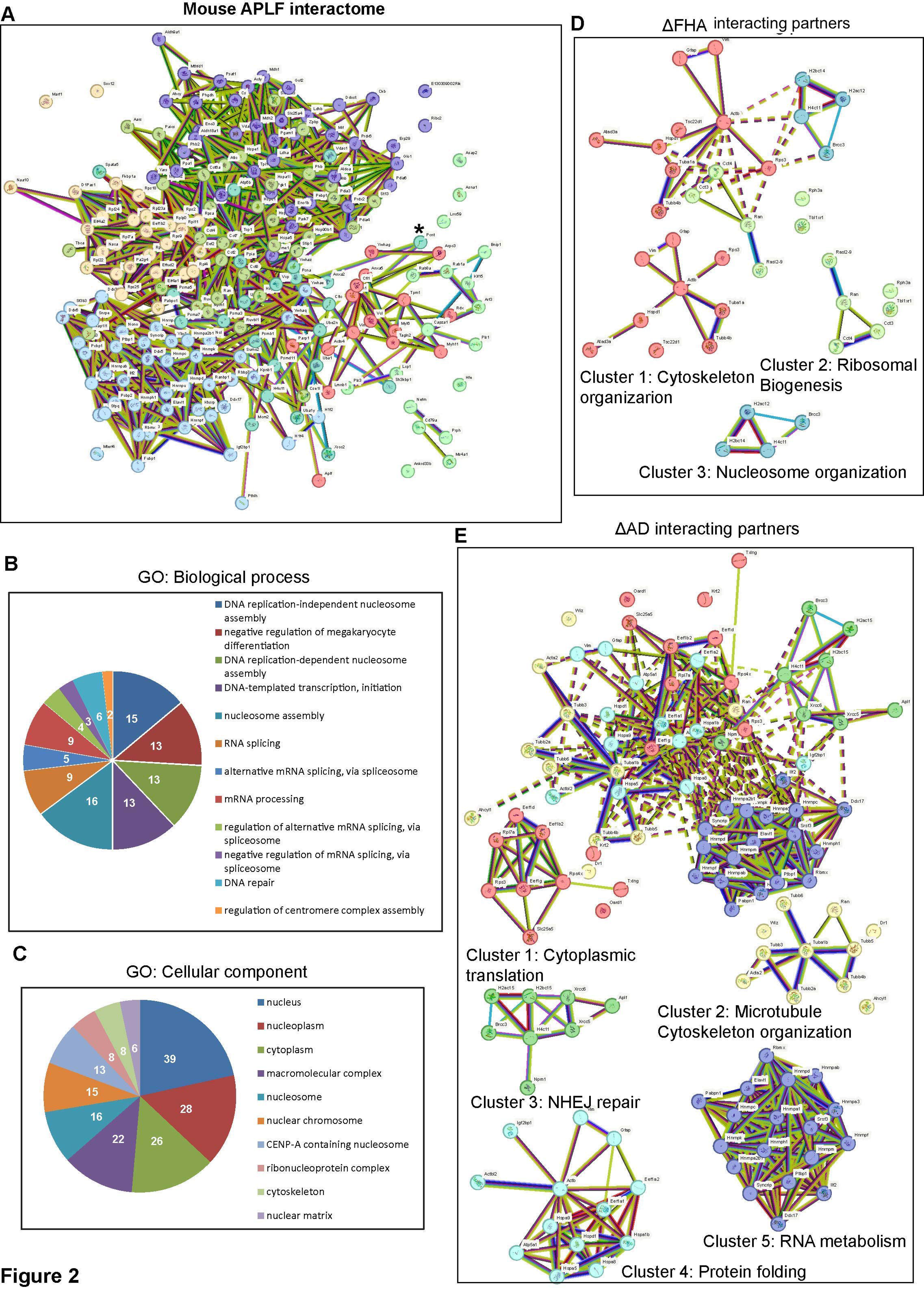
**Mouse APLF interactome**. A. Total protein was isolated from HEK293T cells transfected with FLAG-tagged *Aplf* or empty FLAG vector. APLF-interacting proteins were pulled down using FLAG antibody by co-immunoprecipitation and the complex extracted from Protein-A sepharose beads was run on a SDS-PAGE and the gel piece was subjected to LC/MS-MS analysis for the determination of interacting partners of APLF. STRING database analysis for the APLF interacting partners resulted in the creation of seven distinct clusters. * Indicate PCNT as one of the interacting partners (the relevance has been discussed in the later section of the study). B, C. Pie-chart representing the Gene ontology (GO) analysis for top 10 biological processes and cellular components using DAVID bioinformatics tool (https://david.ncifcrf.gov). D, E. APLF domain specific interaction partners- Total protein was isolated from HEK283T cells transfected with FLAG-tagged ΔAD-APLF, ΔFHA-APLF or empty FLAG vector. Domain specific interacting proteins were pulled down using FLAG antibody by co-immunoprecipitation and the complex extracted from Protein-A sepharose beads was run on a SDS-PAGE and the gel piece was subjected to LC/MS-MS analysis for the determination of interacting partners of different domains of APLF. STRING database analysis for the domain specific APLF interacting partners resulted in the creation of three distinct clusters for ΔFHA domains (D) and three for ΔAD (5).

These findings instigated us to look into the cellular localization of APLF in mouse embryonic stem cells (ESCs).

### APLF localizes to centrosome

Immunofluorescence analysis demonstrated the presence of APLF at the peri-nuclear region of the ESCs (Fig. 3A). Co-localization of APLF with Centrin 2 (CETN2, a centrosomal marker) and pericentrin (PCNT, a pericentriolar material marker) further substantiated its spatial distribution within the centrosome region (Fig. 3B, S3B). It is interesting to note here that, LS/MS-MS study on APLF interactome revealed interaction of APLF with PCNT (Fig. 2A, *marked). However, not all the cells within a single field showed a similar expression pattern. So, does the localization vary with cell cycle phases? Therefore, the ESCs were synchronized at different phases of cell cycle (Fig. S3C). Cells in the interphase showed co-localization of APLF with CETN2 in the S-phase and partial co-localization with γ-Tubulin (TUBG), a centrosomal microtubule marker, and CETN2 in G1/S-phase in ESCs (Fig. 3C, S3D). On the other hand, during all the stages of mitotic cell division, APLF significantly co-localized with TUBG (Fig. 3C). Quantitative analysis indicated significant increase in Pearson correlation coefficient between APLF and TUBG in mitosis in comparison to interphase (Fig. 3D). Upon consideration of co-localization of CETN2 and TUBG with APLF in different phases of interphase and mitosis respectively, a significant increase in Pearson correlation coefficient was observed in mitosis (Fig. 3E). The centrosomal localization of APLF was further confirmed with APLF antibody, antibody #2, procured from another company (Fig. S3E). But, does APLF only localize within the centrosome? Then how can it chaperone histones? It could be noted here that in response to DNA damage induced by etoposide in mouse ESCs, APLF was significantly present within the nucleus, confirmed with both the antibodies of APLF procured from two different sources (Fig. S3F). Our earlier studies also demonstrated that with increase in level of APLF, it localizes within the nucleus in the HEK cells or MDAMB-231 breast cancer cell (Majumder et al., 2018) and at different stages of mouse embryonic development (Varghese et al., 2021).

**Figure 3.**
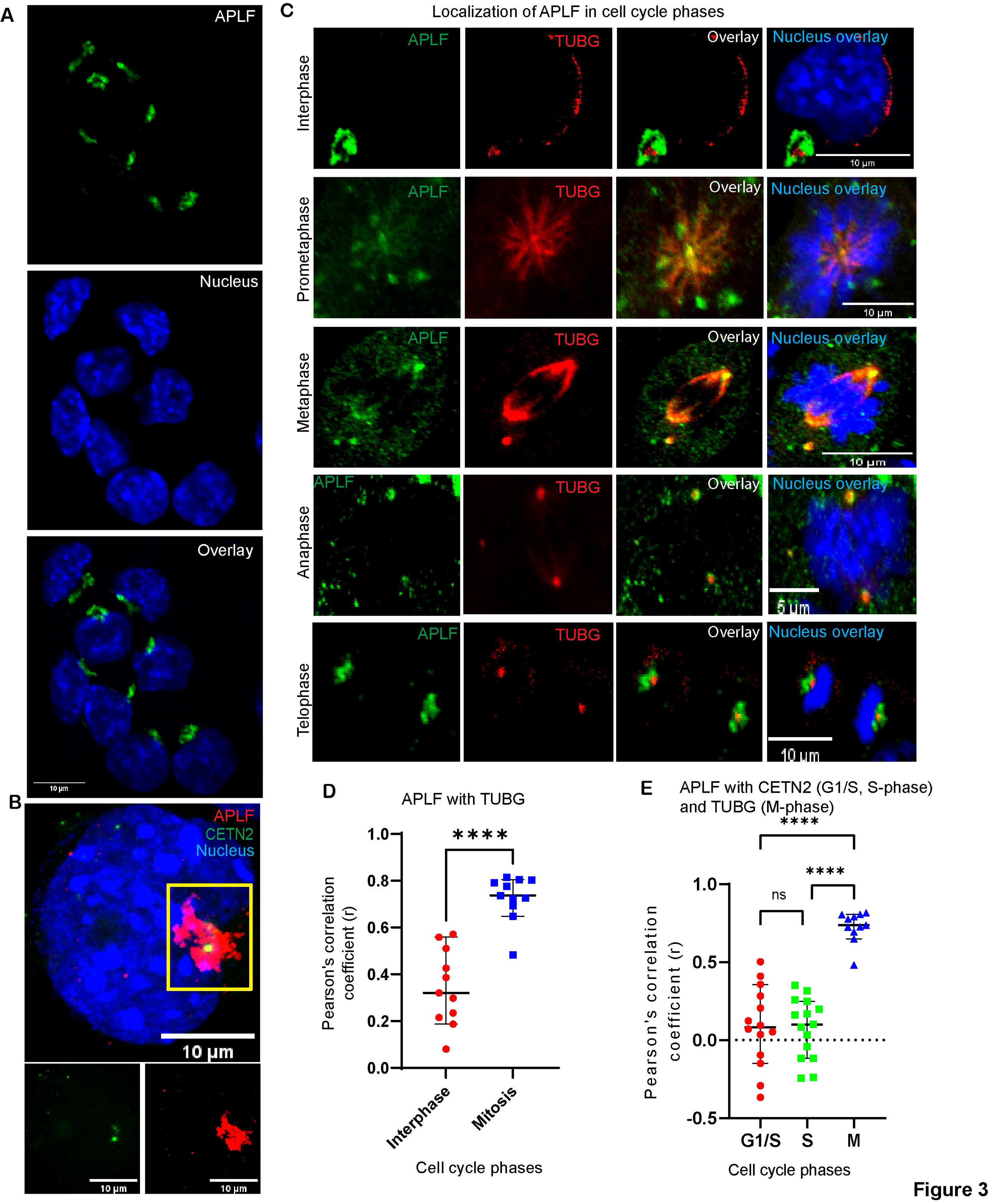
**APLF localizes to centrosomes**. A, B. Immunofluorescence analysis for the expression of APLF in mouse ESCs. B. Immunofluorescence analysis for the co-localization of APLF with CETN2 in mouse ESCs. Inset- APLF and CETN2 expression and co-localization. C. ESCs were synchronized in interphase and mitotic phase (refer to Fig. S3C) of cell cycle under different treatment conditions. These cells were further analyzed for the expression of APLF along with TUBG by immunofluorescence. D, E. Pearson correlation coefficient was determined for APLF co-localizing with TUBG or CETN2 across different cell cycle phases. Ordinary One- way ANOVA was used for statistical analysis followed by Brown-Forsythe test and Tukey’s multiple comparisons tests, ****p<0.0001.

So, does APLF regulate the centrosomes?

### APLF regulate centrosome number

To understand the function of APLF, we stably transfected ESCs with empty FLAG-vector and *Aplf*-FLAG construct and the increase in expression of APLF was confirmed by western blot analysis (Fig. 4A). Next, we detected whether APLF physically interacts with CETN2. Co-immunoprecipitation analysis confirmed the interaction of APLF with CETN2 (Fig. 4B). The centrosome duplication initiates at the G1/S phase. The differential spatial distribution of APLF in interphase and mitosis primed us to address the expression pattern of APLF across cell cycle phases. We generated ESCs stably expressing cell cycle indicator, FUCCI (Fluorescent ubiquitination-based cell cycle indicator) and sorted the cells in G1, G1S and G2M phases for determining the relative expression of *Aplf* (Fig. S4A). *Aplf* was significantly enhanced at the G1/S phase (Fig. 4C) in comparison to other phases of the cell cycle that coincides with the onset of centrosome duplication. Cells synchronized in different cell cycle phases (Fig. S3C) exhibited increased APLF protein expression in the G1/S phase (Fig. 4D). Having observed the centrosomal localization and the peak in expression at G1/S phase, we investigated whether APLF regulates centrosome duplication. Interestingly, the centrosome quantification in APLF over expressing cells, generated in Fig, 4A, revealed an increase in centrosome number in comparison to cells transfected with the empty vector (Fig. 4E, 4F). This increase in centrosome number or supernumerary centrosome upon increased APLF expression corroborated with increase in expression of CETN2 and TUBG (Fig. 4G, S4B). The percentage of cells having ≥3 centrosomes were significantly increased in APLF overexpressing ESCs (Fig. 4H) whereas cells having ≤2 centrosomes were significantly reduced in these cells compared to ESCs with empty FLAG vector (Fig. 4I). In consequent to supernumerary centrosomes, the cells undergoing mitosis demonstrated mitotic aberrations upon overexpression of APLF (Fig. 4J). Quantitative analysis revealed significant increase in number of abnormal multipolar spindle than that of normal or bipolar spindle with increase in level of APLF in ESCs (Fig. 4K). Interestingly, upon downregulation of APLF by shRNA mediated lentiviral transfection (Fig. S4C), we observed an opposite trend in the centrosome number (Fig. S4D). The percentage of cells having ≥3 centrosomes were significantly reduced in APLF knockdown ESCs (Fig. S4D) whereas cells having ≤2 centrosomes were significantly increased in these cells compared to ESCs with scramble shRNA (Fig. S4D). However, we could not observe any significant change in spindle polarity in the *Aplf-*sh ESCs (Fig. S4E).

**Figure 4.**
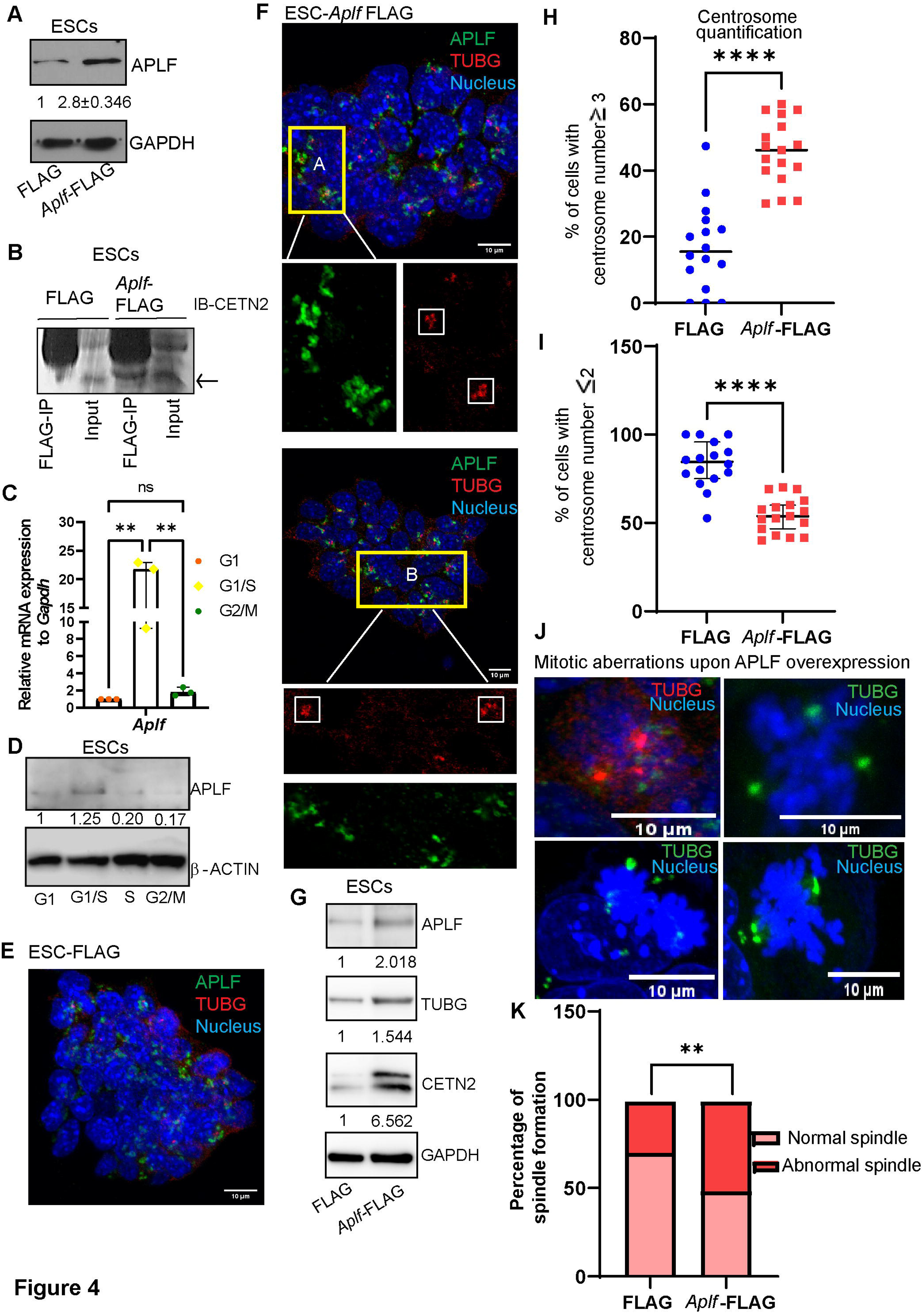
**APLF regulate centrosome number**. A. Western blot analysis for the expression of FLAG-tagged APLF in ESCs. Protein was isolated in RIPA buffer from ESCs stably transfected with the FLAG-tagged mouse *Aplf* and empty FLAG vector, run on SDS-PAGE and probed with APLF antibody. B. Co-immunoprecipitation of APLF and CETN2. Proteins isolated from the above sets were immunoprecipitated with FLAG antibody followed by immunoblotting with CETN2. IgG was used as the negative control. C. ESCs stably expressing FUCCI construct were sorted on the basis of reciprocal expression of CDT1 and its inhibitor Geminin. CDT1 (red) peaks in the G1 whereas the Geminin (green) peaks at G2M. Double positive represents the G1S population (refer to Fig. S4A). Total RNA was isolated from the cell samples and subjected to the generation of cDNA. qRT-PCR analysis was performed to determine the expression of *Aplf* mRNA in different phases of cell cycle. Ordinary one-way ANOVA with Tukey’s multiple comparisons tests were performed for the statistical analysis, **p<0.01. D. Western blot analysis for the expression of APLF in different cell cycle phases synchronized cells (refer to S3C). E, F. Immunofluorescence analysis for the expression of APLF and TUBG in FLAG-tagged constructs in ESCs. ESC-*Aplf* FLAG cells demonstrate increased centrosome number. Inset A and B from the ESC clump shows cells with increased centrosome numbers in different colonies of ESCs (F). G. Western blot analysis for the expression of centrosome associated markers including CETN2 and TUBG in ESCs. H. Scatter plot representing the quantification of number of cells having ≥3 centrosomes in vector control and *Aplf*-FLAG ESCs. I. Scatter plot representing the quantification of number of cells having ≤2 centrosomes in vector control and *Aplf*-FLAG expressing ESCs. A two-tailed unpaired t-test was performed for the statistical analysis, ****p<0.0001. J. Mitotic aberrations formed upon increased expression of APLF in ESCs stably transfected with *Aplf*-FLAG construct, were analyzed with TUBG immunostaining. K. Stacked bars representing the percentage of normal (bipolar) vs. abnormal (multipolar) spindle formation in control FLAG and *Aplf*-FLAG expressing ESCs. One sided Fisher’s exact test was used for the statistical analysis, **p<0.01.

But, how does APLF regulate the centrosome number?

### APLF upstream to master regulator of centrosome duplication PLK4

Polo-like Kinase 4 (PLK4) serves as the primary regulatory factor in the process of centrosome duplication. As cells expressing increased level of APLF resulted in supernumerary centrosome, we were intrigued to understand whether APLF could regulate PLK4 level or not. Hence, we determined the expression of PLK4 in ESCs stably transfected with *Aplf*-FLAG (Fig, 4A). We failed to observe any significant alteration in the level of PLK4 both at the mRNA and protein level (Fig. 5A, 5B). Next we checked whether APLF physically interact with PLK4 or not. Co-IP analysis demonstrated endogenous APLF interact with PLK4 indeed (Fig. 5C, 5D). Phosphorylated PLK4 or activation of PLK4 regulates centrosome duplication. Hence, we looked into the phosphorylation status of PLK4. As mouse specific p-PLK4 antibodies are commercially not available, to my knowledge, firstly we used an indirect approach wherein we immunoprecipitated the ESC derived lysate with pan phospho-Ser/Thr antibody followed by immunoblotting with PLK4 antibody. We observed an increased level of the phospho-PLK4 protein in ESCs expressing *Aplf*-FLAG in comparison to cells expressing empty FLAG vector (Fig. 5E, 5F). Next, determined the p-PLK4 level in ESCs using antibody raised against the p-S305 of human PLK4. This site is conserved in mouse APLF. We observed an increase in the phosphorylation of PLK4 at the S305 site (Fig. 5G). Thus, increase in APLF level corresponded to increase in phosphorylated PLK4 in ESCs.

**Figure 5.**
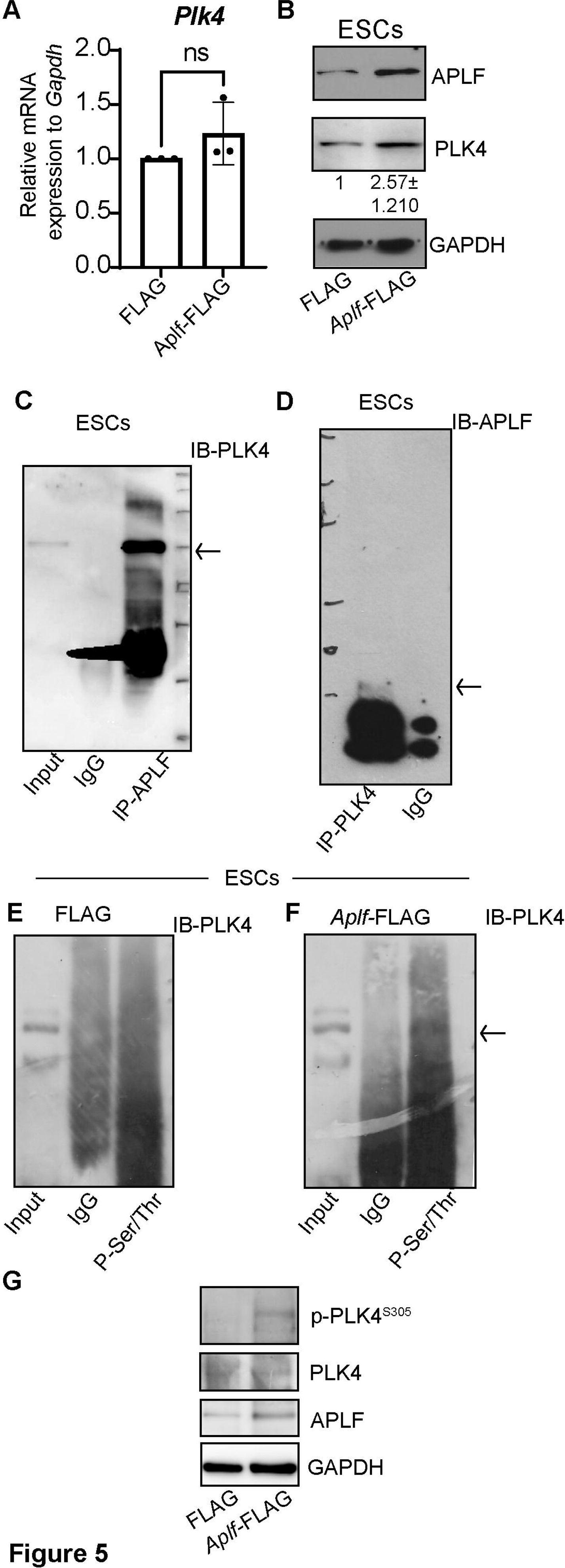
APLF in regulation of PLK4. A. qRT-PCR analysis for the expression of *Plk4* in stably expressing FLAG and *Aplf*-FLAG ESCs. Two-tailed paired t-test was used for the statistical analysis, ns- not significant. B. Western blot analysis for the expression of PLK4 in the same set of cells analyzed in A. C. Co-immunoprecipitation of endogenous APLF with PLK4. Proteins were isolated from ESCs and immunoprecipitated with APLF antibody followed by immunoblotting with PLK4. D. Proteins isolated from ESCs were immunoprecipitated with PLK4 antibody followed by immunoblotting with APLF. E. In the same set of cells analyzed in Fig. 4, proteins were isolated and immunoprecipitated with pan p-Ser/Thr antibody followed by immunoblotting with PLK4. The presence of band at 100KDa confirms that the increased expression of APLF enhanced phosphorylation of PLK4 in ESCs. F. Western blot analysis for the expression of p-PLK4^S305^ in FLAG and *Aplf*-FLAG expressing ESCs. Level of total PLK4, APLF and GAPDH was also determined in the same set of samples.

But, how does increase in APLF level result in increase in phosphorylation of PLK4? Does this indicate any additional characteristic of mouse APLF?

### APLF is a kinase

Phosphorylation of a protein is one of the post-translational modifications that could regulate biological processes and is mediated by the kinase enzyme. Hydrolysis of ATP drives the kinase reaction. To further explore on this context, we first predicted the mouse APLF structure (Fig. 6A) using bioinformatics tool, as crystal structure of the entire molecule is not available till date. The full structure of mouse APLF was modelled using the Iterative Threading ASSEmbly Refinement (I-Tasser) program (Yang and Zhang, 2015). Because of lack of sufficient templates for modeling the whole APLF structure, only FHA domain of mouse APLF was modeled using crystal structure of FHA domain of human APLF (PDB ID: 5W7W) as a template using Modeller 10.1 software (Webb and Sali, 2016). To detect the kinase activity of APLF, we looked for ATP docking site in the APLF molecule. Docking analysis showed presence of 3 sites, D11, R37, K95, within the FHA domain of APLF which might be responsible for the binding of the ATP molecule (Fig. 6B). For the enzymatic analysis, we generated His-tagged APLF clones (Fig. S5A, S5B). Purified full length APLF and domain specific clones from HEK293T cells were used for kinase reaction using ADP-Glo kinase reagent. This assay analyzes ADP produced during kinase reactions. Luciferase reactions generate light via ADP-ATP conversion. The Relative Luminescence Unit (RLU) indicates kinase activity. With increasing amount of APLF, the RLU increased and also with increase in ATP concentration, the substrate saturation curve followed the Michelis-Menten equation for ATP (Fig. 6C, 6D). The K_m_ for ATP= 27.15μM. Hence, the affinity of APLF for ATP is considerably high. Since, FHA domain harbor the ATP docking sites, we looked into the ΔFHA clone to determine the kinase activity (Fig. S5C, S5D). We observed almost complete abrogation of the kinase activity in absence of the FHA domain (Fig. 6E). The empty His-tagged construct was also transfected and his-tagged protein lysate was purified and used as control (Fig. 5E). Furthermore, we generated mutants for all the 3 putative ATP docking sites by site directed mutagenesis and checked for the kinase activity of the mutated APLF clones (Fig. S5E, S5F). We observed significant loss of kinase activity for all the 3 mutants, however, for R37A construct, the kinase activity was entirely abrogated (Fig. 6F). We infer that R37 is essential for the kinase activity. If the kinase activity of APLF is responsible for the increase in centrosome number, then mutation in the R37 residue should also reflect an alteration in the formation of supernumerary centrosomes. Hence, we quantified the centrosome number in ESCs stably transfected with empty vector, *Aplf*-His and APLF^R37A^-His constructs. Cells expressing APLF with mutated R37 exhibited a significant decrease in the population of cells with a centrosome count of three or more, while there was an significant increase in the population of cells with a centrosome count of two or less, when compared to cells expressing the unaltered full-length APLF protein (Fig. 6H, S5G). The cells that expressed the empty vector or APLF^R37A^-His exhibited comparable centrosome numbers, with no statistically significant difference observed between these two groups of cells (Figures 6H, S5G). On the other hand, intact APLF expression resulted in supernumerary centrosome in ESCs, replicating our earlier observation (Fig. 6H, S5G, 4H, 4I). Thus, R37 residue or the kinase activity of APLF is significantly important to regulate the centrosome number in mouse ESCs. The next question is whether APLF is really phosphorylating substrates during the enzymatic reaction or not. To answer that question, we determined the presence of phosphorylated proteins within the product of kinase reaction for APLF at different concentration (Fig. 6C) and hence the samples were run on SDS PAGE and the membrane was probed with pan phospho-Ser/Thr antibody. Interestingly the intensity of a band at ∼100KDa increased with increase in amount of enzyme (Fig. 6I, pointed with an arrow).

**Figure 6.**
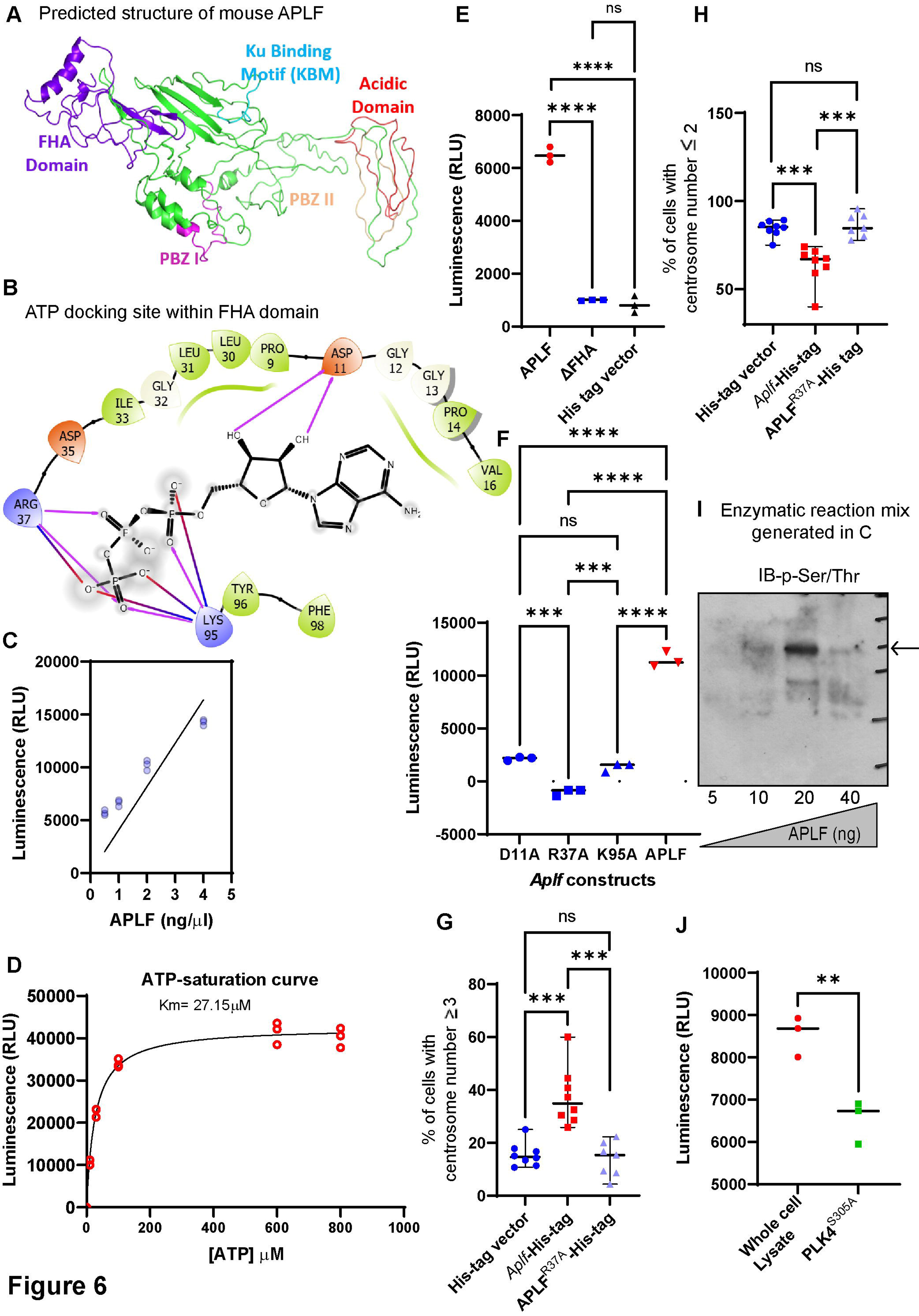
**DNA repair factor and histone chaperone APLF is a kinase**. A. Mouse APLF structure was predicted using the bioinformatics tool, I-TASSER program. B. Docking studies revealed presence of three putative ATP binding sites within the FHA domain. C-K. Detection of kinase activity using ADP-Glo kinase reagent. Different constructs of APLF were cloned into His-tag vector (refer to Fig. S5). Eluate #5 (having 250mM imidazole) was used as the source for the enzyme. C. Graph shows the enzymatic activity of APLF in Relative Luminescence Unit (RLU) with increasing amount of the enzyme APLF. His-tagged clones were transfected in HEK293T cells and the protein was isolated and purified using Ni-NTA agarose beads and eluted in 250mM imidazole. Simple linear regression analysis was performed for the curve fitting. D. The affinity towards ATP was determined by assaying the kinase activity at different concentrations of ATP. Binding of ATP within APLF follows the Michaelis–Menten equation with a K_m_ for ATP= 27.15 μM. Michaelis-Menten least square non-linear fit was used to derive the K_m_ for ATP. E. Scatter plot shows the enzymatic activity of control APLF and ΔFHA His-tagged and His-tagged empty vector control proteins were purified in a similar manner mentioned above. Two-sided unpaired t-test was performed for the statistical analysis, ns- not significant, ****p<0.0001. F. The three ATP binding sites within the FHA domain were mutated by site directed mutagenesis and the His-tagged clones were transfected in HEK293T cells and processed for the isolation of purified proteins. These APLF mutated enzymes were analyzed for the enzymatic activity. Scatter plot shows the loss in enzymatic activity upon incorporation of mutation within the FHA domains. G, H. Control His-tag vector, APLF-His-tag and APLF^R37A^-His- tag constructs were transfected in ESCs for the detection of alteration in centrosome numbers. Immunofluorescence analysis was performed for the expression of TUBG in the cells (refer to Fig. S5G). Scatter plot representing the quantification of number of cells having ≥3 or ≤2 centrosomes in control, *Aplf*-His, APLF^R37A^-His expressing ESCs. Two-sided unpaired t-test was performed for the statistical analysis, ns- not significant, ***p<0.001. I. Western blot analysis for the presence of phosphorylated proteins in the enzymatic reaction mix mentioned in C. The product formed was run on SDS-PAGE and immunoblotted with pan phospho-Ser/Thr antibody. A band at ∼100KDa showed increase in intensity with increasing amount of APLF. J. Scatter plot for the enzymatic activity of APLF in presence of either whole cell lysate or PLK4 mutated at the autophosphorylation site of S305 as the substrate. Two-tailed unpaired t-test performed for the statistical analysis, **p<0.01.

It can be recalled that the size of PLK4 is ∼100KDa. In addition to this, intensity of few more protein bands increased upon increase in amount of APLF in the kinase reaction. This indicates that APLF might have multiple substrates for kinase activity. In order to confirm whether PLK4 is a substrate for APLF, we isolated and purified 3XFLAG-tagged PLK4^S305A^ (Addgene #69839) (Sillibourne et al., 2010) protein from HEK293T cells and used as substrate for the kinase reaction. The S305 residue of PLK4 is associated with its autophosphorylation and its role in centrosome duplication. The present construct has S substituted with A amino acid. We observed that there is significant loss in the kinase activity of APLF in presence of S305 mutated PLK4 substrate; however, it is not completely abolished (Fig. 6J). We predict two possibilities from this outcome. Firstly, APLF might phosphorylate PLK4 at other sites in addition to S305 or APLF might phosphorylate other substrates in addition to PLK4.

Hence we provide experimental evidence in delineating kinase activity of APLF in regulation of centrosome number in mouse ESCs (Fig. 7) independent of its histone chaperone activity.

**Figure 7.**
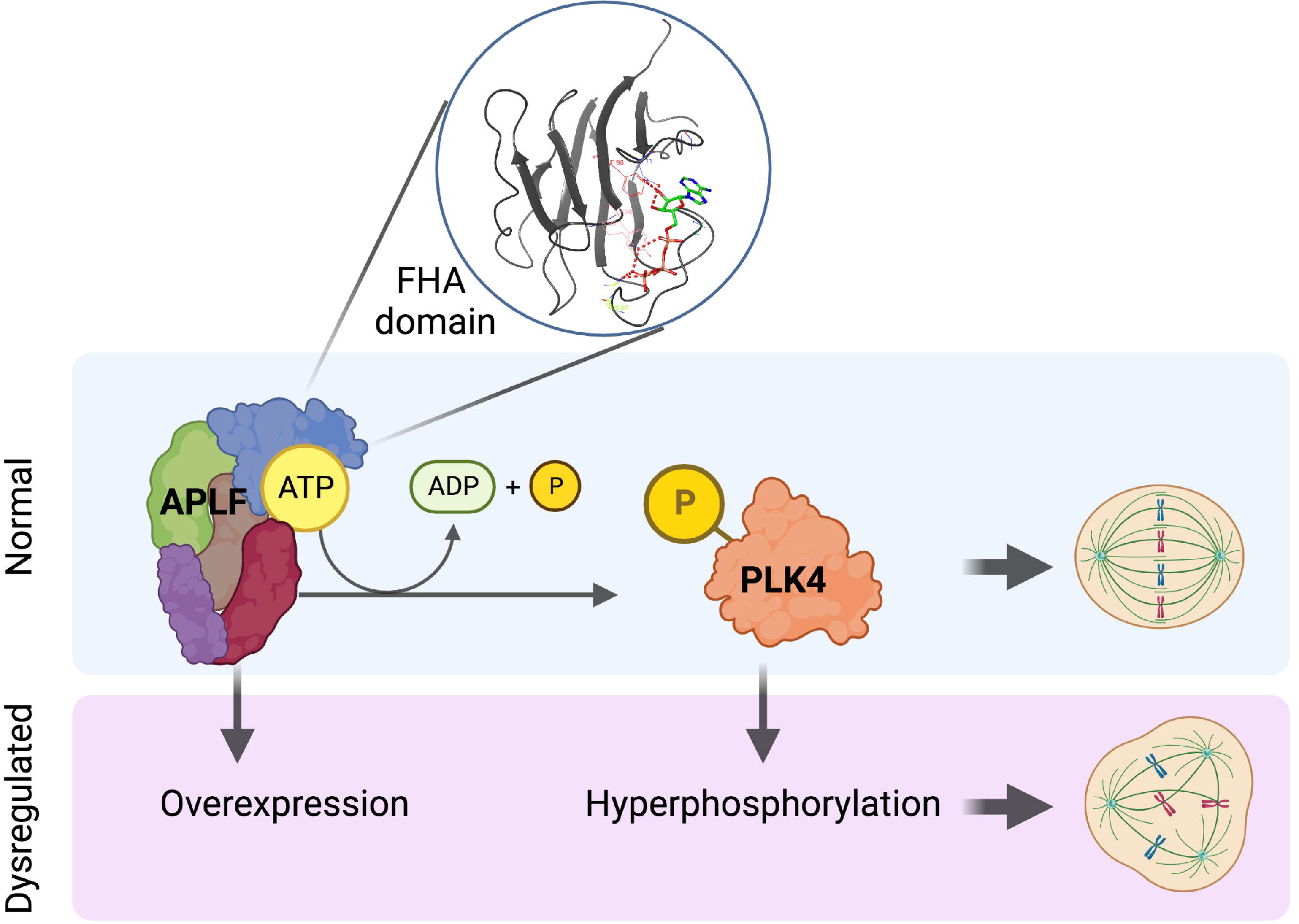
Model depicting APLF as kinase in regulation of centrosome number-. The kinase activity of APLF regulate PLK4 phosphorylation thereby regulating centrosome number. The model has been generated with Biorender.

## Discussion

Rapidly dividing cells are likely to have a high rate of chromosomal aberrations and compromised genetic stability. Replication error, impairment in spindle orientation, and chromosome mis-segregation contribute to genetic instability in rapidly dividing cells like stem cells. The centrosome plays a crucial role as a prominent organizer of microtubules and serves as a pivotal spindle pole throughout the process of mitosis. It is composed of a pair of centrioles and a protein matrix known as pericentriolar material. The potential consequence of deregulating the process of centriole assembly is the occurrence of centrosome amplification, which in turn can result in aneuploidy. Aberrant centrosome structure and number causes cancer and developmental defects and ciliopathies. The centrosome serves as a primary sequestration site for the DNA repair factors, cell cycle regulators, majorly the checkpoint proteins in mouse ESCs (Koledova et al., 2010). PLK4 is the master regulator of centriole assembly, centrosome amplification and is the major kinase driving centriole duplication (Coelho et al., 2015; Ryniawec et al., 2023). On the other hand, depletion or blockage of kinase activity of PLK4 inhibits centriole duplication (Wong et al., 2015). While it was shown that trans-autoactivation of PLK4 initiates centriole biogenesis (Lopes et al., 2015) but its autophosphorylation at S305 residue results in centriole duplication (Sillibourne et al., 2010). However, the stretch of 24 amino acids from 282-305, called the degron, is required for the (in)stability of the active PLK4 which regulates the centrosome number (Holland et al., 2010; Cunha-Ferreira et al., 2013). But, studies have suggested that other kinases might be present in the system which can phosphorylate PLK4 (Lopes et al., 2015). Our study gives an indication that APLF could be that kinase which can phosphorylate PLK4 at both S305 and other residues such that its activation results in supernumerary centrosomes. However, we are yet to decipher which exact site of PLK4 is phosphorylated.

Although we could demonstrate that APLF lies upstream to PLK4 thereby regulating centriole duplication, but its exact position in the cartwheel structure is yet to be revealed along with its role in the regulation of proteins, namely, PLK1, SASS6, CEP152, STIL among others. Our results showed the increased expression of both CETN2 and TUBG upon increased expression of APLF (Fig. 4G) and this kind of report has already been observed in case of increase in number of centrosome (Sillibourne et al., 2010). This further consolidates the role of APLF in dictating centrosome number. Our earlier studies in murine development demonstrated loss in implantation sites around E7.5 (Varghese et. al., 2021), and interestingly, PLK4 knockout causes embryonic arrest at E7.5 embryonic stage (Hudson et al., 2001). While direct experimental evidence is lacking, it is plausible to hypothesize that the inactivation of PLK4 due to loss of APLF may impede developmental processes. Our initial experiments showed that in mouse embryo injected with *Aplf*-overexpressing constructs, led to a delay in the pre- implantation development (data not shown) in comparison to early abnormal hatching in presence of *Aplf-*knockdown constructs during pre-implantation development (Varghese et al., 2021).

Several constraints were encountered throughout the course of this study. Initially, the worry regarding the mobility of the aggregates necessitated the execution of the FRAP experiment in HEK293T cells instead of ESCs. In order to establish a comparable experimental context, we employed identical cell samples to assess the DNA repair capabilities of the APLF domains and as well as the interactome studies. Nevertheless, this approach turned out to be advantageous since the expression level of APLF in HEK293T cells is significantly lower when compared to ESCs. Previous research from our lab has already substantiated this discovery (Syed et al., 2016; Majumder et al., 2018). The lack of a commercially accessible phospho-PLK4 antibody with reactivity towards the mouse species was a significant obstacle. Therefore, we determined the p-PLK4 both by immunoprecipitation analysis followed by p-PLK4 antibody specific for human p-PLK4^S305^. Determining the precise phosphorylation location of PLK4 would provide a comprehensive understanding of the regulatory mechanism employed by APLF in relation to PLK4. Nevertheless, previous research has indicated that the identification of the precise phosphorylation location within the dregon poses considerable challenges. However, employing site-directed mutagenesis on individual sites within the dregon could potentially facilitate the identification of the precise phosphorylation site for APLF within PLK4.

It is commonly seen as advantageous to produce the desired mammalian protein within a prokaryotic system in order to minimize any ambiguity arising from the presence of other proteins, whether as contaminants or related factors, that may interfere with the protein of interest. However, we choose to refrain from utilizing that technique in order to specifically isolate the protein in its active state. There is a significant possibility that the protein may lose its activity when expressed in the prokaryotic environment. In order to mitigate the presence of contaminants or proteins that may be bound, we employed an empty His-tag vector for the purpose of analysis. Moreover, the utilization of deletion domain or mutants has yielded functional evidence regarding the isolation of APLF.

Till date, only one histone chaperone, Nucleophosmin, has been identified as a substrate of Cyclin E/CDK2 for initiating centrosome duplication (Okuda et al., 2000; Wang et al., 2005), while no other histone chaperone has been shown to demonstrate a kinase activity. Hence, APLF stands out in both these aspects. As the ATP docking site lies within the FHA domain, it also remains to be seen whether the particular sites have any influence on the DNA repair activity or not. FHA domains are binding modules that recognize only phospho-Thr residues. These domains are present in both prokaryotic and eukaryotic proteins that play significant roles in different cellular phenomena including transcription, DNA damage-repair, cell cycle regulation (Reinhardt and Yaffe, 2013). However, the association of a kinase activity of this domain has never been reported. Hence, we anticipate that this finding might open up unforeseen characteristics of multiple proteins associated with the FHA domains.

The observation that despite a mere 59% homology in amino acid sequences, the domains in mice and humans exhibit conservation, serves to enhance the significance of exploring APLF. Hence, we predict that human APLF might also possess a kinase activity. The utilization of human ESCs in clinical settings is hindered to a great extent by the presence of genomic instability, which is partly attributed to the occurrence of supernumerary centrosomes during the process of mitosis in the culture environment (Holubcová et al., 2011). The role of centrosomes has been well documented in stem cell division and differentiation (Schatten and Sun, 2011). Therefore, the kinase activity of APLF might contribute to a better understanding for the maintenance of ESCs in undifferentiated or differentiated states that could potentially influence their usage or application in different regenerative purposes.

## Materials and Methods

### Cell culture and synchronization

HEK293T was cultured in standard DMEM (Himedia #AL0078) containing 10% FBS (Invitrogen #1600044) and 1% Penicillin/ Streptomycin (Invitrogen #15140122) and 1% Antimycotic/Antibiotic (Invitrogen #15240062), as described previously (Majumder et al., 2015). E14 ESCs were cultured in feeder-free conditions in ES culture media containing IMDM (Invitrogen #12440053; Himedia #AL070A), 15% ESC qualified serum (Invitrogen #10439024), 0.0124% monothioglycerol (Sigma #M6145), 1% Penicillin/ Streptomycin, 1% Antimycotic/Antibiotic and Leukemia Inhibitory Factor (LIF-10^5^U/ml) (Millipore #ESG1107) (Syed et al., 2016).

Double thymidine treatment was given for synchronization in G1/S phase as described previously (Chen and Ding, 2018). E14 ESCs were treated with 2mM Thymidine (Sigma #T9250) for 18 hours followed by the release by washes with pre-warmed PBS and recover in fresh media for 6 hours. Then second thymidine (2mM) treatment was given for 18 hours and collected 0hour and 7.5hours for G1S and G2M respectively. For S phase synchronization, the cells were treated with 50µg/ml Hydroxyurea (Sigma #H8627) for 18hours.

### Plasmid constructs and Site-Directed Mutagenesis

Mouse APLF full-length was cloned as previously described (Majumder et al., 2018). The cDNA of APLF full length and APLF^ΔFHA^ and APLF^ΔAD^ were amplified with respective primer pairs (Table S1) and subcloned into Flag-HA-pCDNA3.1 (Addgene #52535) (Horn et al., 2014), pcDNA^TM^3.1/V5-His A (a kind gift from, Dr. K B Harikumar, RGCB), pEGFP-N1 (a kind gift from Dr. Arumugam Rajavelu, RGCB).

The ATP binding mutants (APLF^D11A^, APLF^R37A^, APLF^K95A^) were generated using Q5® Site-Directed Mutagenesis Kit (NEB #E0554) according to the manufacturer’s protocol. The mammalian expression vector pcDNA^TM^3.1/V5-His A was used to construct the His-tagged ATP binding mutants. The primers have been listed in Table S1.

### Stable transfection of mouse APLF

The mouse APLF full-length, the deletion domain constructs (APLF^ΔFHA^ and APLF^ΔAD^) and the mutated APLF^R37A^ along with their respective vector controls (Table S1) were ectopically expressed in E14 ESCs using lipofectamine 2000 reagent (Invitrogen #11668-019), according to the earlier reports (Majumder et al., 2018). The transfected cells were selected for 14 days with 250μg/ml G418, a Neomycin analog (Sigma #G8168) and used for further analysis.

### Protein extraction and immunoblotting

The total cell lysate was prepared by lysing the cells by sonication in radioimmunoprecipitation assay (RIPA) buffer (10mM Tris-HCl, pH7.5, 150Mm NaCl, 5mM EDTA, 1% Triton X-100, 1% Nonidet P-40, 1% Sodium deoxycholate, 0.1%SDS, 1mM Sodium orthovanadate, and 1mM PMSF) (Majumder et al., 2015). Protein concentration was estimated using Bradford Reagent (Bio-Rad #500-0006). An equal amount of protein extracts was loaded for different samples for immunoblotting in 8%, 10%, or 12% resolving SDS-PAGE. Specific antibodies (Table S2) were probed after transferring the protein to the PVDF membrane and visualized by enhanced chemiluminescence.

### Co-immunoprecipitation

The total cell lysate was prepared by multiple freeze-thaw cycles followed by sonication in NETN100 buffer (20mM Tris-HCl, pH7.5, 100mM NaCl, 1mM EDTA, 0.1% Nonidet P-40, 1mM Sodium orthovanadate, 1mM Sodium fluoride, protease inhibitor cocktail-1tablet) and incubated overnight with the antibody of our interest (Majumder et al., 2015). The antibody-lysate complex was incubated with protein A- conjugated sepharose beads (Sigma #P3391) and eluted in 2X laemelli dye, resolved by SDS-PAGE, and analyzed by immunoblotting.

### Quantitative real-time PCR (qRT-PCR)

Total RNA was extracted using Qiagen RNAeasy Kit (Qiagen #74106) according to the manufacturer’s protocol. cDNA was generated with random oligomer primers by using cDNA reverse transcription kit (ABI Biosystems #4368814). qRT reaction was carried out using and Sybr Green master mix (ABI Biosystems #4309155) (Varghese et al., 2021). The qRT-PCR primers used are listed in Table S3.

### *In vivo* nucleosome assay using Fluorescence recovery after photobleaching (FRAP)

HEK293T cells were co-transfected with H2B: mCherry and GFP tagged APLF deletion domain constructs (APLF, APLFΔAD, APLFΔFHA) using calcium phosphate method in 8-well chambered Lab-Tek coverslips. Cells were then synchronized in S phase after 24 hours of transfection before the bleaching experiments. FRAP experiments were performed on a Zeiss LSM 980 with Airyscan-2 confocal light scanning microscope (Carl Zeiss, Germany), equipped with 40X/0.95 NA, non-oil objective. To photo-bleach the H2B: mCherry, we selected a region of interest (ROI) of 5µm 2 in the nucleus and bleached with 561nm laser and the fluorescence recovery observed at the intervals of 10s over a time-course of 3minutes. The unbleached ROI in the nucleus and background ROI of 5µm 2 were also recorded while photobleaching. Two images were captured before bleaching. Bleaching intensity and scanning time set to obtain the minimum bleaching efficiency of 50% in shortest possible time. The FRAP analysis was done using ZEN Blue software. The fluorescence intensity of bleached, unbleached, and background ROI was obtained, and the background correction was done. The fluorescence recovery of the bleached nucleus was quantified relative to the unbleached nucleus. The fluorescence intensities were also normalized with mean prebleach intensity and expressed in recovery percentage. The recovery curve was plotted with normalized fluorescence intensity over time, and the curve fitting was done with a one-phase association equation for linear regression in GraphPad prism. The half-time of recovery and the mobile fraction correspond to the mean plateau value, and halftime was estimated from the recovery curve.

### DNA damage induction and repair assay

The mouse APLF full-length, the domain deletion constructs (APLF, APLF^ΔAD^, APLF^ΔFHA^) transfected ES E14 cells were treated with 10μM etoposide (Sigma #E1383) for 4hours. Subsequently, the treatment was removed and cells were thoroughly washed with pre-warmed PBS and allowed the damage to recover for different time. The cells collected at different time points (0h, 6h, 24h) were analyzed by immunostaining for the presence of γH2A.X (Abcam #ab2893) foci (Syed et al., 2016).

### Immunofluorescence and Imaging

Immunofluorescence analysis was conducted as per earlier protocol (Syed et al., 2016). The cells grown on 18mm or 14mm coverslips were fixed with 100% methanol and blocked in 1% BSA. Cells were subsequently incubated with primary antibodies (Table S2) overnight followed by the labeling with secondary antibodies conjugated with Alexafluor 488/568. Nuclear staining was performed with Hoechst dye 33258 (1 µg/ml) and the cells were mounted in Prolong Antifade (Invitrogen #P36934). Images were acquired in Olympus FV3000 and Nikon A1Rsi Confocal Laser Scanning Microscope equipped with 60Xobjective, oil immersion. Serial optical sections along Z-axis were obtained by the sequential scan of 0.3μm step size and merged to obtain the maximum-intensity projection. Image analysis and centrosome quantification were performed using Fiji (ImageJ) software.

### Docking Methodology

Because of lack of proper templates for modeling the whole APLF structure, only FHA domain of mouse APLF was modeled using crystal structure of FHA domain of human APLF (PDB ID: 5W7W) as a template using Modeller 10.1 software . To understand ATP interaction with mouse APLF FHA domain, we subsequently docked ATP with APLF FHA domain using Autodock 4.2.6. Autodock thoroughly searches ligand conformations and assesses the free energy of binding to the target. It employs an amber force field and a free energy scoring algorithm based on linear regression analysis. Using AutoDockTools, Gasteiger charges were applied to all atoms, and rotatable bonds were assigned. The binding energies of proteins and ligands were determined using atom affinity potentials precalculated on grid maps using AutoGrid. The affinity grid maps centered on the protein were 120×120×120 Å in size, with 0.375 grid point spacing. Using the Lamarckian method, Autodock 4.2.6 calculates the ligand binding across the conformational search space. The final docked conformations were clustered using a tolerance of 1 Å root-mean-square deviation (RMSD).

### Protein expression and purification

The His-tagged mouse APLF full-length, the domain deletion constructs (APLF, APLF^ΔAD^, APLF^ΔFHA^), and ATP binding mutants (APLF^D11A^, APLF^R37A^, APLF^K95A^) were expressed and purified from a mammalian cell line, HEK293T. The cells were lysed by multiple cycles of freeze-thawing followed by sonication with extraction buffer (20mM Tris-HCl, pH7.5, 200mM NaCl, 10ug/ml PMSF, 1μg/ml Leupeptin). The lysate was incubated with equilibrated Ni-NTA Agarose resins (Qiagen #30210) at 4ℒC for 2-4 hours. The resin-bound His-tagged proteins were washed with wash buffer containing 20mM imidazole (Sigma #12399) to eliminate non-specific binding and eluted 5 times subsequently with elution buffer (20mM Tris HCl pH7.5, 200mM NaCl, 250mM imidazole). The purified protein was run on SDS-PAGE to determine the purity of the isolated protein by Coomassie staining.

### LC/MS-MS analysis

In order to identify the prospective interaction partners, ESCs transfected with APLF full length and domain deletion constructs were lysed with NETN100 lysis buffer described previously (Baral et al., 2023). The lysate was precipitated with FLAG (Abcam #ab1162) antibody and the pull-down samples were subjected to proteomic analysis by liquid chromatography tandem mass spectrometry Synapt G2 High Definition MS™ System (HDMS E System) (Waters, Manchester, UK) attached to nanoACQUITY UPLC ® chromatographic system (Waters, Manchester, UK) at the proteomic facility, RGCB, India. Data were analyzed and the peptide hits were annotated using DAVID bioinformatics Resources (Sherman et al., 2022) and STRING database. Proteins showing at least one unique peptide have been considered.

### *In vitro* kinase assay

*In vitro* kinase assays were performed as previously described (Jain et al., 2019). 200ng of heat-inactivated ES cell lysate (substrate) were mixed with different concentrations (10ng, 20ng, 30ng, 40ng) of purified APLF constructs in kinase buffer (25mM Tris-HCl pH7.5, 2mM DTT, 10mM MgCl2, 0.1mM Sodium orthovanadate, and 10μM ATP) and incubated at 30°C for 1hour. Kinase reactions were stopped with ADP-Glo^TM^ Reagent (Promega #V6930) followed by the addition of ADP Glo^TM^ Kinase detection reagent (Promega #V6930) to convert ADP to ATP and introduce luciferase to detect ATP. Luminescence was detected using Varioskan^TM^ Flash multimode reader (Thermo Scientific). A standard curve for ATP was prepared with differing concentrations of ATP according to the manufacturer’s protocol.

## Acknowledgements

We thank Dr. Suparna Sengupta (RGCB), Dr. Ananda Mukherjee (RGCB) and Dr. Tapas Manna (IISER, Thiruvananthapuram) for their valuable inputs and sharing reagents. We thank the RGCB Mass-spectrometry and Confocal facilities. The work is partially supported by Science & Engineering Research Board (SERB), Department of Science & Technology (#CRG/2022/005438, #CRG/2021/002528), ICMR (2021-9968/SCR/ADHOC-BMS), and intramural fund from the institute aided by Department of Biotechnology. SMR, MBS received fellowship from DST INSPIRE and UGC respectively (#IF180251 and #191620091128). KN received funding from SERB (SRG/2022/001697).

## Author contributions

SMR performed experiments, analyzed data, prepared figures, along with DD wrote manuscript; PCV performed experiments, generated stable ESC expressing *Aplf*-FLAG ESCs; MBS designed cloning primers and generated the SDM clones; SKM generated APLF domain constructs tagged with FLAG and checked their expression in HEK293T cells; AD performed the DNA repair assay; KN deduced the APLF structure and the docking studies; DD conceived the idea, performed experiments, analyzed data, along with SMR wrote the manuscript and with permission from all the authors submitted the manuscript.

## Abbreviation

AD: Acidic Domain
APLF: Aprataxin PNK-Like Factor
CDK2: Cyclin Dependent Kinase 2
CETN2: Centrin 2
DSB: Double Strand Break
EMT: Epithelial-Mesenchymal Transition
ESCs: embryonic stem cells
FHA: Forkhead Association Domain
FRAP: Fluorescence Recovery After Photobleaching
FUCCI: Fluorescent Ubiquitination-based Cell Cycle Indicator
GO: Gene ontology
KBM: Ku Binding Motif
NHEJ: Non-Homologous End Joining
NPM: Nucleophosmin
PBZ: Poly (ADP Ribose) Binding Zinc Finger
PLK4: Polo-like Kinase 4
RAN: Ras-related Nuclear protein
ROI: Region of interest
SDM: Site-directed Mutagenesis
TUBG: γ-Tubulin

