## Supplementary information for "Kinase activity of histone chaperone APLF maintains steady state of centrosomes in mouse embryonic stem cells"

**Supplementary methods**

**RNA interference-** Mouse *Aplf* shRNA validated from our earlier study (Syed et al., 2016) was used to knockdown *Aplf* in ESCs. Lentiviral supernatant was produced in HEK293T cells by transient transfection using calcium chloride. Briefly HEK293T cells were grown to 80% confluency. The plasmids pRRE (gag/pol), pMD2G (VSVG), pRSV (Rev), and *Aplf* shRNA were combined in a solution containing 0.25M $CaCl_2$ and then balanced with an equivalent amount of 2× HEPES-buffered saline. The supernatants containing lentiviral particles were collected following a 24-hour incubation period. ESCs were grown to 70% confluence and transfected with lentiviral soups. Transfected cells were selected by the addition of 1 μg/ml of puromycin (Sigma, #P8833). After 3 days, protein was extracted for analysis. Western blotting confirmed the knockdown.

**Figures**

Colour-coded for amino acid conservation

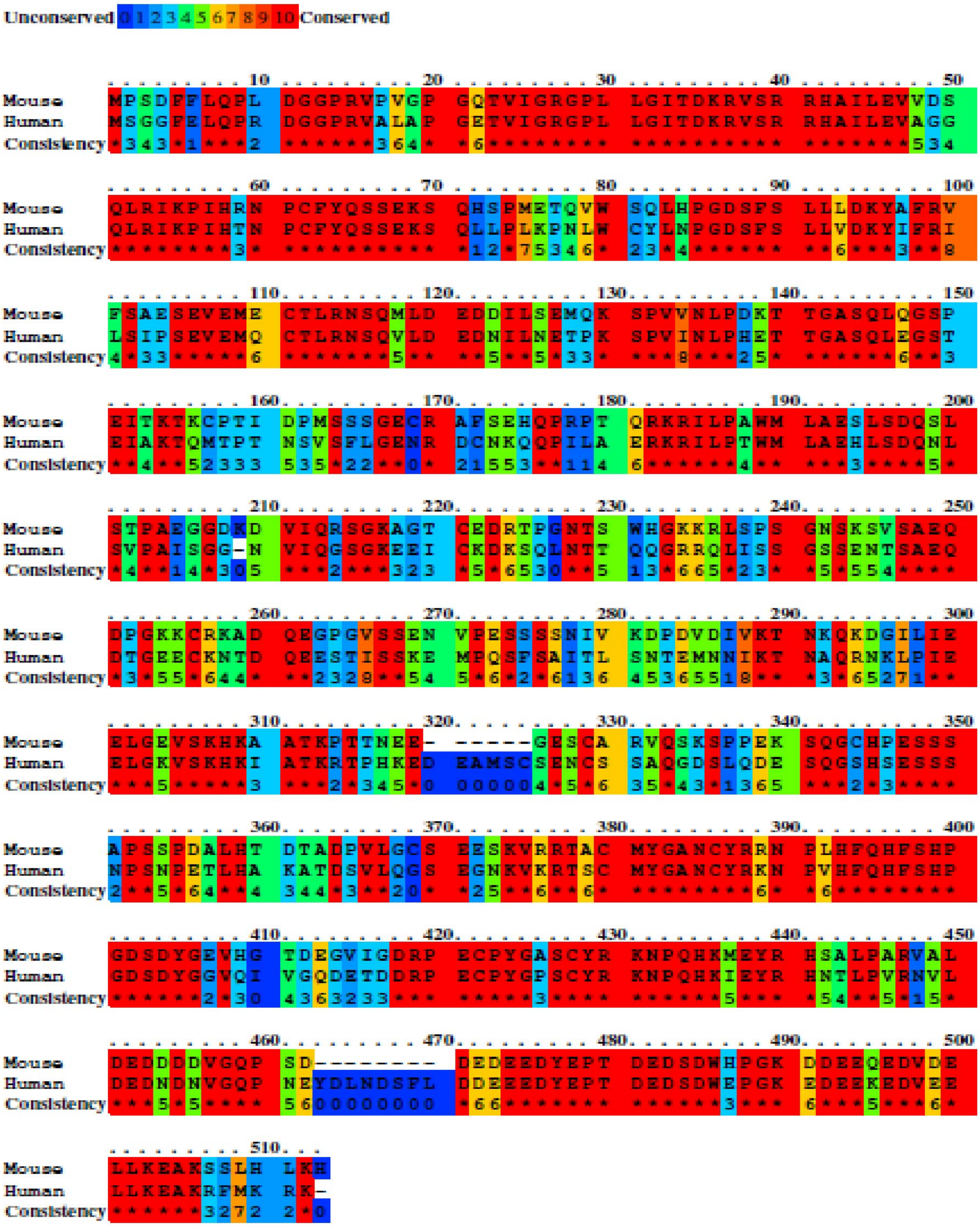


Figure S1

**Figure S1.** Amino acid conservation of mouse and human APLF sequences. FHA, Ku binding, PBZ and AD domains marked within the APLF protein sequence.

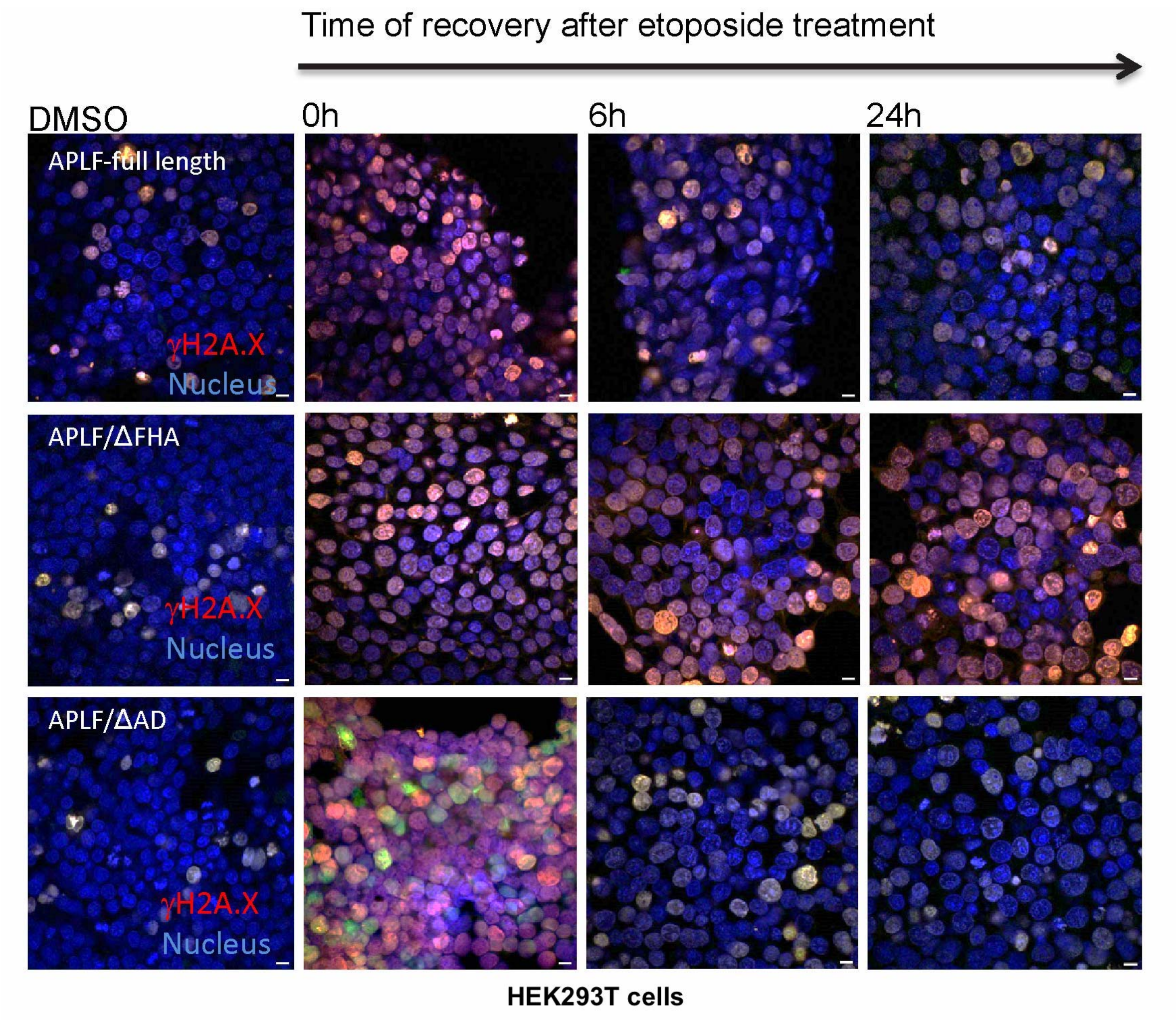


### Figure S2

**Figure S2.** DNA repair activity of APLF domain. HEK293T cells were treated with either 10μM etoposide or DMSO for 4 hours, followed by recovery till 24 hours. Presence of γH2AX foci was determined as mark of DNA damage by immunofluorescence analysis corresponding to different domain constructs of APLF.

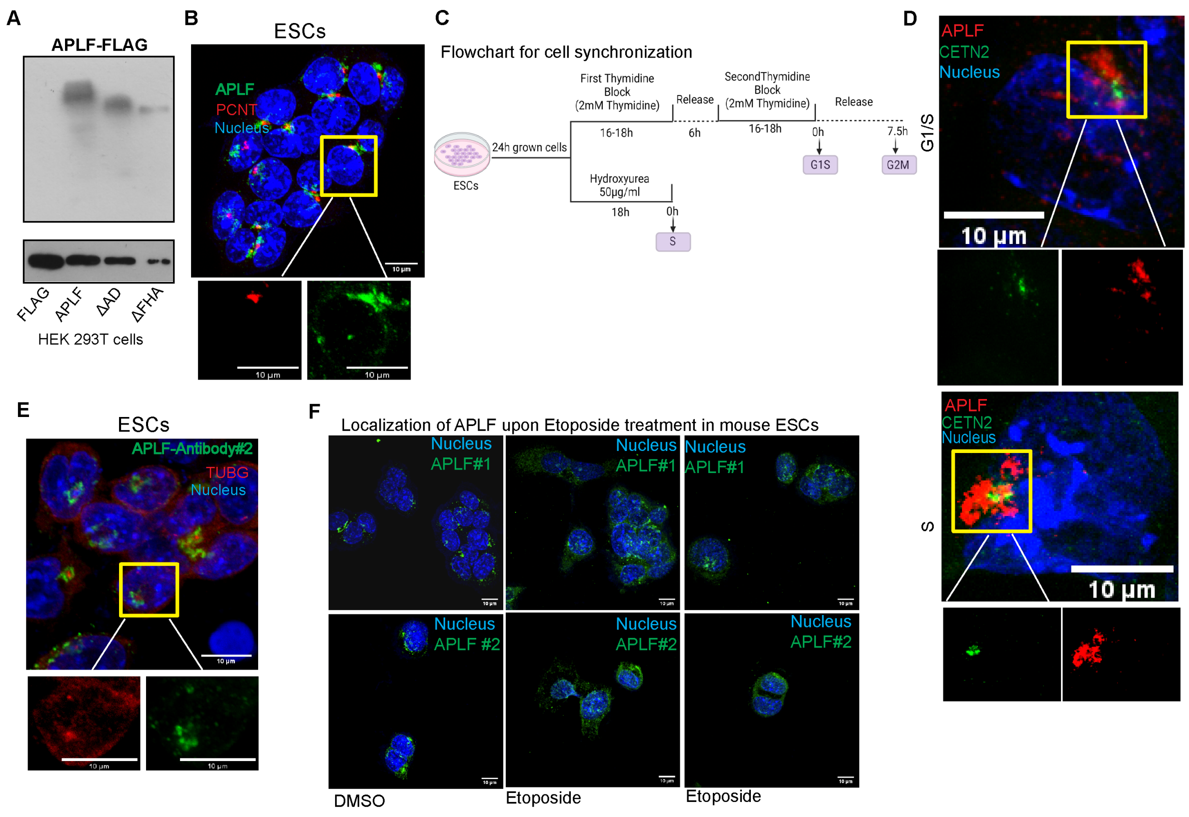


**Figure S3**

**Figure S3**. A. Western blot analysis for the expression of APLF-FLAG constructs. The constructs were transfected in the HEK293T cells followed by whole cell protein isolation in RIPA buffer and run on SDS-PAGE. The resulting gel was probed with FLAG antibody. B. Immunofluorescence analysis for the expression of APLF and PCNT in mouse ESCs. Inset shows the magnified image of TUBG and APLF expression. C. Flow chart representing the ESC synchronization, generated using Biorender. D. Immunufluorescence analysis for the expression of APLF along with CETN2 in cells synchronized at G1 and G1/S phase of the cell cycle. Inset shows the magnified image of TUBG and APLF expression. E. Immunofluorescence analysis for the expression of APLF by additional antibody #2 procured from another company. Inset shows the magnified image of TUBG and APLF expression F. Immunofluorescence analysis for the expression of APLF by antibody #1 and #2 in presence or absence of etoposide (10µM).

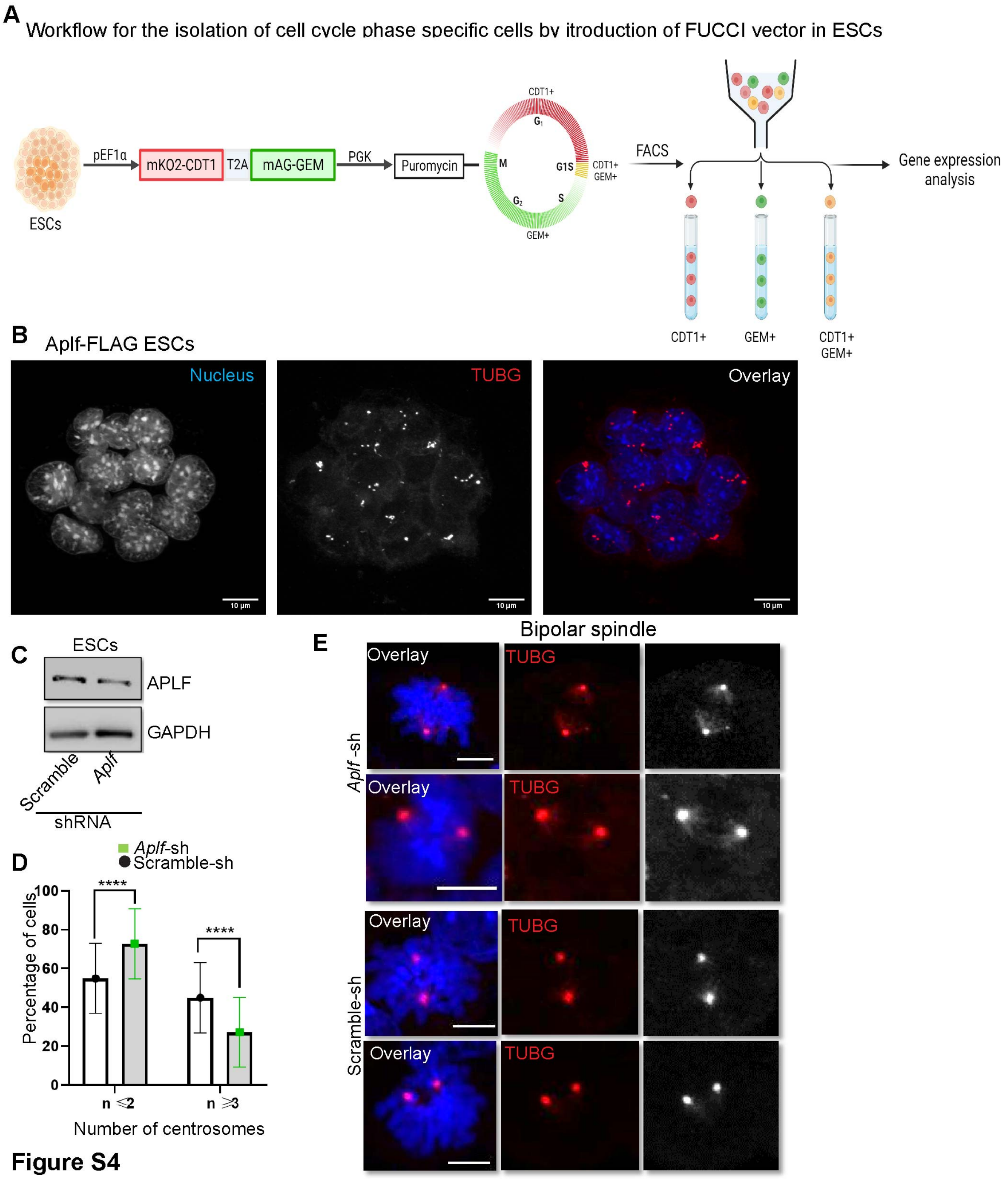


### Figure S4

**Figure S4.** A. Workflow for the generation and usage of FUCCI expressing ESCs. B. Immunofluorescence analysis for the expression of TUBG in *Aplf*-FLAG expressing ESCs. C. Western blot analysis for the expression of APLF in shRNA mediated knockdown of APLF in ESCs in comparison to cells transduced with particles having scramble shRNA. D. Bar graphs represent the number of cells having ≥3 or ≤2 centrosomes in scramble shRNA vs. *Aplf*-sh expressing ESCs. E. Immunofluorescence analysis for the expression of TUBG within the mitotic spindle in Scramble shRNA and *Aplf*-sh expressing ESCs.

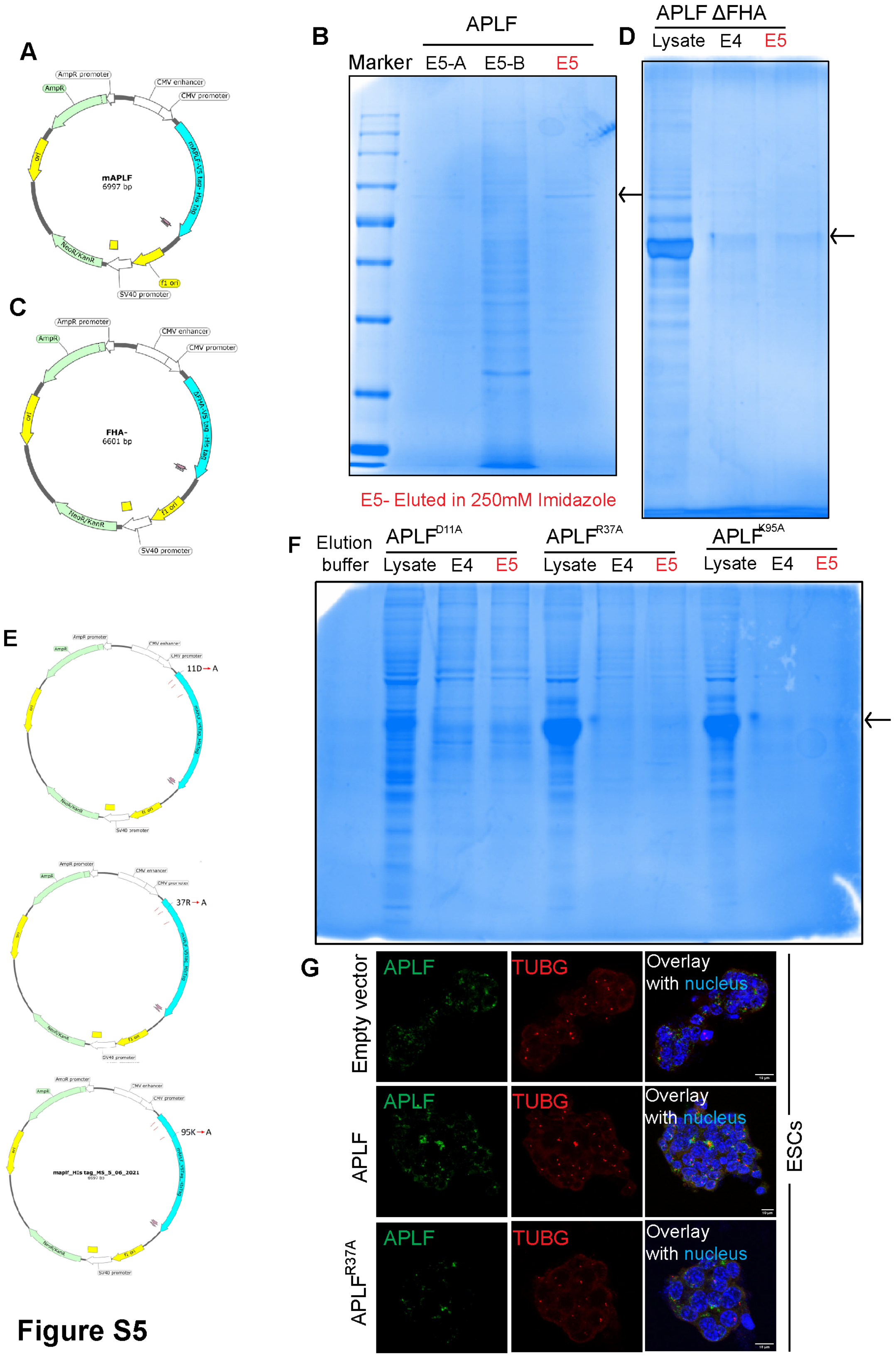


**Figure S5**

**Figure S5.** A. Vector map of *Aplf* ligated to His-tag vector. B. Coomassie stained SDS-PAGE demonstrate the purification *Aplf*-His tagged protein from the transfected HEK293T cells. The eluate E5 mentioned in red font has been used in this study. C. Vector map of

FHA ligated to His-tag. D. Coomassie stained SDS-PAGE demonstrate the purification of FHA-His tagged protein isolated from the transfected HEK293T cells. The eluate E5 mentioned in red font has been used in this study. E. Vector map of *Aplf* mutants ligated to His-tag. F. Coomassie stained SDS-PAGE demonstrate the purification His-tagged mutant proteins isolated from the transfected HEK293T cells. The eluate E5 mentioned in red font has been used in this study. G. Immunofluorescence analysis for the expression of APLF and TUBG in ESCs transfected with empty His-vector, *Aplf*-His and $APLF^{R37A}$-His.

**Table S1: List of cloning primers used in the study**

| Primer | Forward | Reverse |
|---|---|---|
| APLF:GFP | GGGGTACCATGCCTTCGGACTTCTTTC | CGGGATCCCCCTTTTTCCTTCTCATAAAC |
| APLFΔAD:GFP | GGGGATCCCCCAGAGCAACTCTGGCTGG | GGGGTACCATGCCTTCGGACTTCTTTC |
| APLFΔFHA:GFP | GGGGTACCATGGAGTGTACTCTGAGAAAC | CGGGATCCCCCTTTTTCCTTCTCATAAAC |
| APLF:His | CGTAGGTACCATGCCTTCGGACTTCTT | TGCTCTAGACTTTTTCCTTCTCATAAACC |
| APLFΔAD:His | CGTAGGTACCATGCCTTCGGACTTCTT | TGCTCTAGACAGAGCAACTCTGGCTG |
| APLFΔFHA:His | CGTGGTACCTGTACTCTGAGAAACAGTCAG | TGCTCTAGACTTTTTCCTTCTCATAAACC |
| APLF$^{D11A}$:His | CAGCCTCTGGCCGGCGGTCCC | CAGAAAGAAGTCCGAAGGCATGGTACC |
| APLF$^{R37}$A:His | AACAGACAAAGCAGTATCCAGAAGACATGCCATTCTTGAAGTG | ATTCCCAGCAGCGGCCCG |
| APLF$^{K95A}$:His | GTTACTTGACGCGTACGCTTTCCGTG | AGGGAAAAGCTATCCCCG |
| APLF:FLAG-HA | GGCTCTAGAATGCCTTCGGACTTCTTT | GGCGGATCCTCAATGTTTCAAGTGCAA |

**Table S2: List of antibodies used in the study**

| Antibodies | Dilution (Applications) | Company (catalog #) |
|---|---|---|
| APLF | 1:2000 (IB); 1:200(IF)<br>1:100 (IP) | Invitrogen (PA539776) (antibody #1), Sigma (SAB4500756) (antibody #2) |
| CETN2 | 1:2000 (IB) | Proteintech (15877-1AP) |
| PLK4 | 1:2000 (IB) | Proteintech (1295-1AP) |
| GAPDH | 1:10000 (IB) | Sigma (g9545) |
| β-ACTIN | 1:2000 (IB) | Abcam (ab8226) |
| Phospho Ser/Thr | 1.5ul/0.1μg (IP) | Cell Signaling (9631S) |
| γH2AX | 1:200(IF) | Abcam (ab2893) |
| TUBG | 1:400(IF) | Abcam (ab11316) |
| FLAG | 1:2000(IB) | Abcam (ab1162) |
| GFP | 1:1000(IB) | Cell Signaling (2956) |
| p-PLK4$^{S305}$ | 1:500 (IB) | ThermoFisher (PPLK-140AP) |
| Peroxidase Affinipure Goat Anti Rabbit IgG | 1:7000, 1:10000 (IB) | Jackson Immuno Research (111035144) |
| Goat Anti Mouse IgG-HRP | 1:2000 (IB) | Santacruz (SC2005) |
| Goat Anti-Rabbit Alexaflour 488 | 1:200 (IF) | Invitrogen (A11008) |
| Goat Anti-Mouse Alexaflour 568 | 1:200 (IF) | Invitrogen (A11004) |

**Table S3: List of qRT-PCR primers**

| Gene | Forward | Reveree |
|---|---|---|
| *Aplf* | CAAGGAAGCCCTGAAATAAC | TGAAAGCTCTGCATTCACCT |
| *Plk4* | GGAGAGGATCGAGGACTTTAAGG | CCAGTGTGTATGGACTCAGCTC |
| *Gapdh* | TGCCCCCATGTTTGTGATG | TGTGGTCATGAGCCCTTCC |
